# Differential effects of PD-1 and CTLA-4 blockade on the melanoma-reactive CD8 T cell response

**DOI:** 10.1101/2020.12.15.422827

**Authors:** Anastasia Gangaev, Elisa A. Rozeman, Maartje W. Rohaan, Daisy Philips, Sanne Patiwael, Joost H. van den Berg, Antoni Ribas, Dirk Schadendorf, Bastian Schilling, Ton N. Schumacher, Christian U. Blank, John B.A.G. Haanen, Pia Kvistborg

## Abstract

Immune checkpoint inhibitors targeting programmed cell death protein 1 (PD-1) and cytotoxic T-lymphocyte-associated protein 4 (CTLA-4) have revolutionized the treatment of melanoma patients. Based on early studies addressing the mechanism of action, it was assumed that PD-1 blockade mostly influences T cell responses at the tumor site. However, recent work has demonstrated that PD-1 blockade can influence the T cell compartment in peripheral. blood. If activation of circulating tumor-reactive T cells would form an important mechanism of action of PD-1 blockade, it may be predicted that such blockade would alter either the frequency and/or the breadth of the tumor-reactive CD8 T cell response. To address this question, we analyzed CD8 T cell responses towards 71 melanoma associated epitopes in peripheral blood of 24 melanoma patients. We show that both the frequency and the breadth of the melanoma-reactive CD8 T cell response in peripheral blood was unaltered upon PD-1 blockade. In contrast, a broadening of the melanoma-reactive CD8 T cell response was observed upon CTLA-4 blockade, in concordance with our prior data. On the basis of these results, we conclude that PD-1 and CTLA-4 blockade impact the tumor-reactive CD8 T cell response in a distinct manner. In addition, the data provide an argument in favor of the hypothesis that anti-PD-1 therapy may primarily act at the tumor site.

**One sentence summary:** We demonstrate that, contrary to CTLA-4 blockade, PD-1 blockade in melanoma patients does not lead to an increase in the breadth or magnitude of the melanoma-reactive CD8 T cell response in peripheral blood, thereby arguing for a dominant effect of PD-1 blockade at the tumor site.

## Introduction

Immune checkpoint targeting therapies, in particular those targeting the programmed cell death protein 1/ligand 1 (PD-1/PD-L1) axis, now form the standard of care for advanced melanoma *(1)* and a number of other solid cancers including non-small-cell lung cancer (NSCLC) *(2)*, renal-cell carcinoma *(3)* and urothelial carcinoma *(4)*. In spite of the very widespread clinical use of PD-1/PD-L1 blocking agents, the mechanism by which these therapies enhance immune-mediated tumor control remains incompletely established. Early work addressing the mechanism of action of PD-1 blockade showed increased numbers of intratumoral proliferating (Ki-67+) CD8 T cells and T cell receptor (TCR) clones after treatment *(5)*. These findings provided the first evidence for a boosting effect on tumor-specific CD8 T cells at the tumor site. In line with these findings, PD-L1 expression on tumor cells, which drives T cell exhaustion and contributes to tumor escape, has been shown to have a predictive value for therapy outcome *(5–7)*, and influence the activity of PD-1 blockade in at least some mouse models *(8, 9)*. Other studies, however, have suggested that PD-1 blockade may exert its effect through activation of circulating tumor-specific CD8 T cell responses. First, studies in mouse models of chronic viral infection have shown the recruitment of CXCR5^+^ Tim-3^-^ CD8 T cells from the white pulp of the spleen upon PD-1 blockade *(10, 11)*. Second, data obtained using mouse tumor models demonstrated that the proliferative response to anti-PD-1 therapy is dependent on CD28-mediated co-stimulation *(12)*, and these findings are in line with a mechanistic study showing that PD-1 signaling inhibits T cell functionality through attenuation of CD28 co-stimulation *(13)*. Collectively, these latter findings have been interpreted as indicating that PD-1 blockade may induce proliferation of tumor-specific CD8 T cell pools in lymphoid tissues that subsequently migrate to tumor tissues. Recent data from clinical studies in patients provide evidence supporting such a hypothesis. First, NSCLC and melanoma patients treated with PD-1 blockade showed an increase in proliferating (Ki-67+) CD8 T cell subsets *(14–17)*. Second, PD-1 blockade was shown to result in clonal replacement of tumor-infiltrating CD8 T cells in patients with squamous cell carcinoma *(18)*. In contrast, however, analysis of peripheral blood from melanoma patients showed no consistent increase in TCR diversity after treatment *(19)*. Although the current knowledge suggests that PD-1 blockade may alter the tumor-specific CD8 T cell compartment systemically, direct evidence for such hypothesis is currently lacking.

Importantly, the vast majority of studies that have addressed the impact of PD-1 blockade, focused the analyses on the bulk CD8 T cell pool in peripheral blood without assessing the T cell specificity. However, work from Schreiber and colleagues in mouse models has shown that ‘bystander’ CD8 T cells respond differentially to checkpoint targeting therapies compared to tumor-reactive CD8 T cells *(20)*. These findings demonstrate the importance of dissecting the mechanism of action of checkpoint targeting therapies on the tumor-reactive CD8 T cell response rather than the bulk CD8 T cell compartment. In this study, we assessed whether PD-1 blockade can increase the magnitude of pre-existing melanoma-reactive CD8 T cells (boosting) and induce novel tumor-specific CD8 T cell responses (broadening) in peripheral blood of melanoma patients (Fig. 1A).

**Figure 1:**
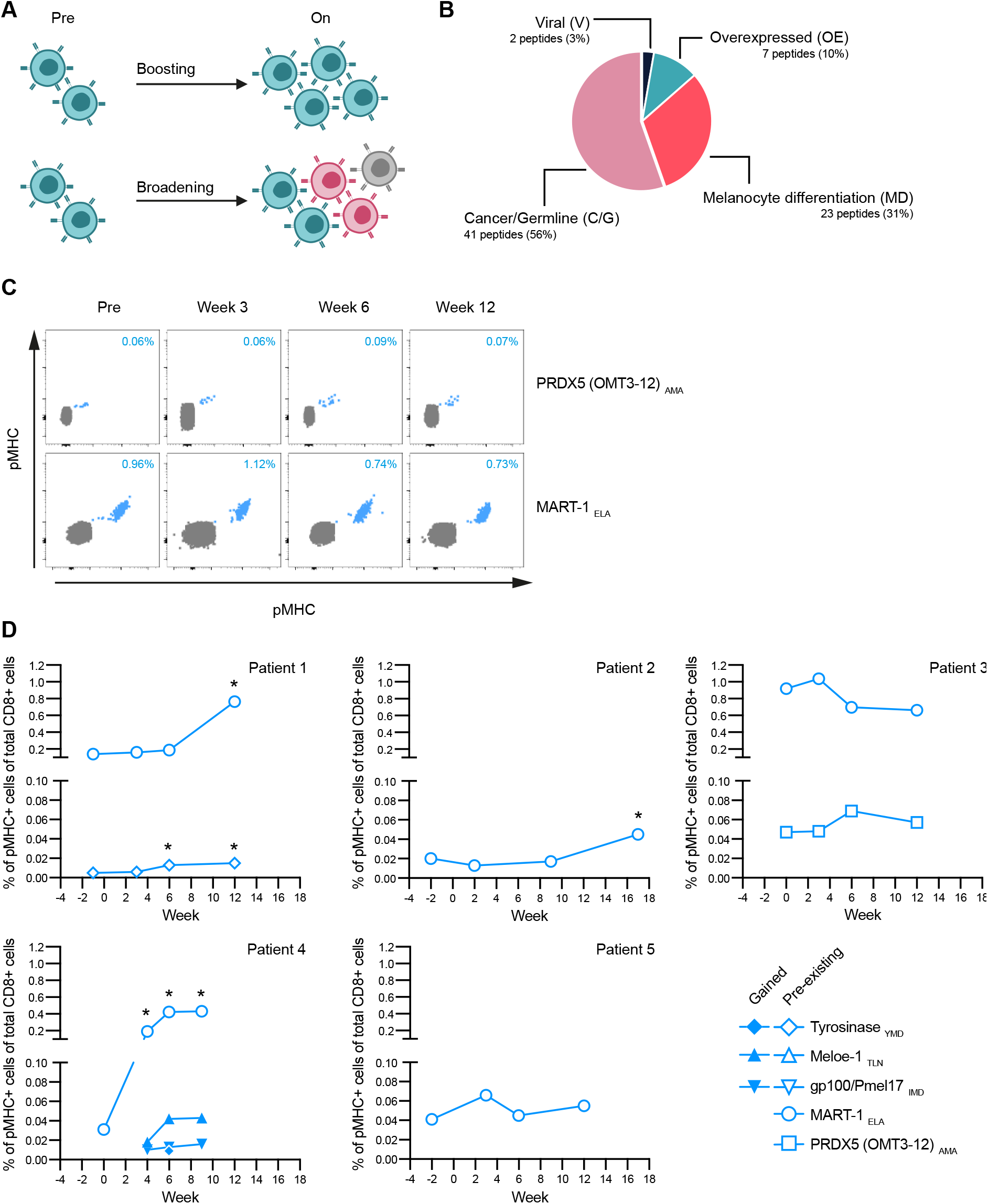
Hypothesis and kinetics of melanoma-reactive CD8 T cell responses during anti-PD-1 therapy. **(A)** Potential mechanisms of anti-PD-1 therapy: expansion of pre-existing tumor-specific CD8 T cells (boosting) and induction of novel tumor-specific CD8 T cell responses (broadening). **(B)** Overview of the HLA-A*02:01-restricted epitope panel. A total of 71 shared melanoma-associated epitopes were included to analyze the tumor-reactive CD8 T cell responses. Two viral epitopes served as positive control for the generation of pMHC multimers. Detailed information is provided in Table S1. **(C)** Representative flow cytometry plots of melanoma-reactive CD8 T cell responses (blue, located in the diagonal of the plot due to the dual coding strategy) pre and on anti-PD-1 therapy. Magnitude of melanoma-reactive CD8 T cell responses (blue, upper right corner) represents the percentage of total CD8 T cells (grey). A representative example of the full gating strategy is provided in Figure S2. pMHC: peptide-major histocompatibility complex. **(D)** Kinetics of melanoma-reactive CD8 T cell responses detected in 5 melanoma patients following anti-PD-1 therapy. A ≥ 2-fold increase (on vs. pre) in magnitude is indicated (*). pMHC: peptide-major histocompatibility complex.

## Results

### Kinetics of the melanoma-reactive CD8 T cell response during PD-1 blockade

To determine which on-therapy time point would allow us to evaluate therapy-induced alterations, we first assessed the kinetics of melanoma-reactive CD8 T cell responses following start of PD-1 blockade. For this purpose, peripheral blood samples from six melanoma patients were collected pre-therapy and at three on-therapy time points between week 3 and week 17 of treatment. The melanoma-reactive CD8 T cell response was analyzed using a panel of peptide major histocompatibility complex (pMHC) multimers loaded with 71 different epitopes derived from previously described shared melanoma antigens restricted to HLA-A*02:01 (Fig. 1B and Table S1). CD8 T cells reactive towards these epitopes were identified by combinatorial encoding of pMHC multimers using a unique dual fluorochrome code for each of the 71 epitopes *(21–23)*. Of note, all responses were confirmed using a different color code combination in each of the two independent experiments. We identified a total of seven melanoma-reactive CD8 T cell responses in five of six patients (Fig. 1C and D). An increase in magnitude (≥ 2-fold) was found in four of the seven identified CD8 T cell responses. In one of the five patients in which melanoma-reactive CD8 T cell responses were detected, three newly detectable (gained) melanoma-reactive CD8 T cell responses were identified. Two of these gained responses were detected at all analyzed on-therapy time points. Together, these data suggested that potential effects of anti-PD-1 therapy in terms of broadening and/or boosting of the melanoma-reactive CD8 T cell response can be detected at the time of the first clinical response evaluation around week 12.

### PD-1 blockade does not alter the melanoma-reactive CD8 T cell response in peripheral blood

To systematically assess potential alterations in terms of boosting and/or broadening of the melanoma-reactive CD8 T cell response upon anti-PD-1 therapy, we included pre- and on-therapy samples from 18 additional patients, resulting in a sample set of 24 patients. For all 48 samples, T cell reactivities towards the 71 melanoma associated epitopes were analyzed, resulting in more than 3400 individual analyses of antigen-specific CD8 T cell responses. Two viral epitopes derived from Epstein-Barr virus and influenza A were included in the analysis as a technical control for the generation of pMHC multimers, and for all panels of pMHC multimers generated virus-specific T cell responses were detected in at least 1 patient (Figure S1). Melanoma-reactive CD8 T cell responses were found in 19 out of 24 patients. The median magnitude of these responses (n=27) was 0.019% of total CD8 T cells (range: 0.005% to 0.920% of total CD8 T cells) (Fig. 2A and B). CD8 T cell reactivity was found towards seven of the 71 melanoma associated epitopes (9%) including epitopes derived from overexpressed antigens (PRDX5 and Meloe-1), cancer/germline antigens (MAGE-A3) and melanocyte-differentiation antigens (gp100, MART-1 and Tyrosinase). Furthermore, in line with prior studies in HLA-A*02:01 positive melanoma patients *(21, 24)*, CD8 T cells reactive towards MART-1 were found in the vast majority of patients (79%). Strikingly, the magnitude of pre-existing melanoma-reactive CD8 T cell responses was unaltered upon therapy with a median fold change of 1.2, demonstrating the lack of a substantial boosting effect (p=0.26) (Fig. 2C). Furthermore, newly detected responses (i.e. responses not detectable before treatment) were only observed in three of the 24 patients (p=0.25) (Fig. 2D). These data show that the frequency and the breadth of the peripheral melanoma-reactive CD8 T cell response was not substantially altered upon PD-1 blockade.

**Figure 2:**
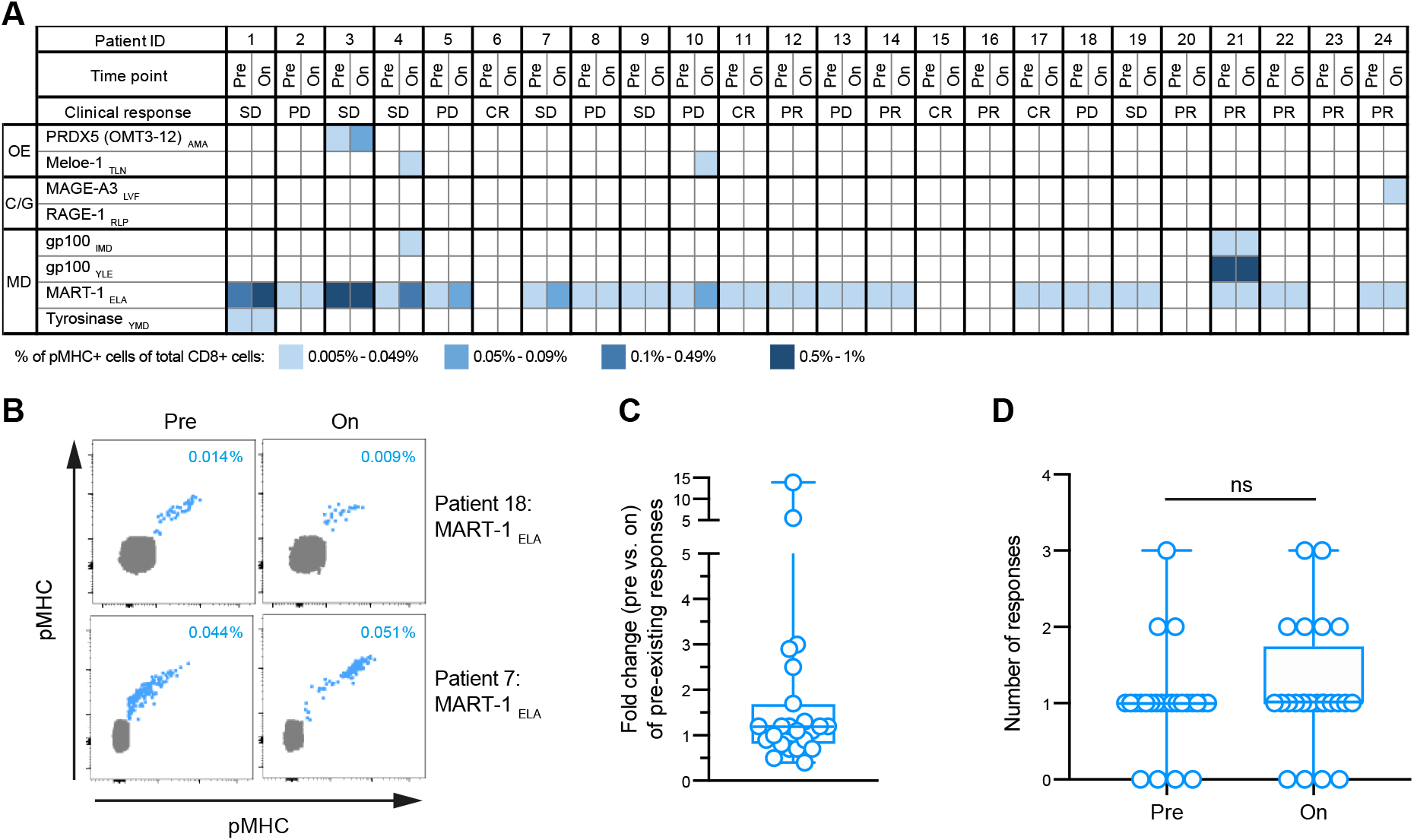
Large-scale analysis of the melanoma-reactive CD8 T cell response upon PD-1 therapy. **(A)** Heatmap overview of melanoma-reactive CD8 T cell responses detected in 24 melanoma patients treated with anti-PD-1 therapy. OE: overexpressed, C/G: cancer/germline, MD: melanocyte differentiation, pMHC: peptide-major histocompatibility complex. **(B)** Representative flow cytometry plots of melanoma-reactive CD8 T cell responses (blue) detected in peripheral blood of two patients pre and on anti-PD-1 therapy. Magnitude of melanoma-reactive CD8 T cell responses is indicated on the top right as the percentage of total CD8 T cells for each response. A representative example of the full gating strategy is provided in Figure S2. pMHC: peptide-major histocompatibility complex. **(C)** Fold change in the magnitude of pre-existing melanoma-reactive CD8 T cell responses (n = 23) after anti-PD-1 therapy. Boxplot represents the median and interquartile ranges, and the whiskers represent the full range. Statistical significance for a change in magnitude on-therapy as compared to pre-therapy was tested with two-tailed Wilcoxon matched-pairs test, p=0.26. **(D)** Number of responses detected in each individual patient (n = 24) pre and on anti-PD-1 therapy. Boxplots represent the median and interquartile ranges, and the whiskers represent the full range. Statistical significance for a change in the number of responses on-therapy as compared to pre-therapy was tested with two-tailed Wilcoxon matched-pairs test, p=0.25.

### PD-1 blockade does not alter the circulating melanoma-reactive and CXCR5^+^ Tim-3^-^ CD8 T cell compartment

Previous studies have shown that anti-PD-1 therapy can induce proliferation of CXCR5^+^ Tim-3^-^ CD8 T cells which are recruited from the white pulp of the spleen in mice with chronic viral infection *(10, 11)*, and similar subsets responsive to anti-PD-1 therapy have been identified in tumor models *(25, 26)*. A change in the frequency of this subset has only been detected early after start of anti-PD-1 therapy (day 7-10 on-therapy) *(16)*. Similarly, data from melanoma patients showed a transient and pronounced proliferation of PD-1^+^ CD8 T cell subsets in peripheral blood at week 1 after treatment initiation *(16)*. Together these findings suggest that PD-1 blockade may induce the priming and recruitment of CD8 T cells from lymphoid organs that can be detected in peripheral blood. To test this hypothesis, we collected peripheral blood samples from eight melanoma patients at week 1, in addition to pre-therapy and week 12. First, we examined alterations in the frequency of the CXCR5^+^ Tim-3^-^ CD8 T cell subset that was previously identified in mice *(10, 11, 25, 26)*. In contrast to chronic viral infection and mouse tumor models, we observed no significant changes in the frequency of CXCR5^+^ Tim-3^-^ CD8 T cells in peripheral blood of melanoma patients, at either week 1 or at week 12 after start of therapy (Fig. 3A and B). Second, we assessed changes of the melanoma-reactive CD8 T cell response in four patients positive for HLA-A*02:01. A total of six melanoma-reactive CD8 T cell responses were found in three of these four patients (Fig. 3C). For two of the three patients no significant alterations in the magnitude of pre-existing melanoma-reactive CD8 T cell responses were found at either week 1 or at week 12. Significant alterations in the magnitude were only detected in one patient (patient 21), in which all three responses showed a transient increase at week 1. Similarly, only one newly induced response was found in one patient (patient 25) at week 1, however, this response was of very low magnitude (0.008% of total CD8) and just above the cutoff (0.005% of total CD8) of the technology. Overall, we observed no consistent boosting or broadening of the melanoma-reactive CD8 T cell response at week 1.

**Figure 3:**
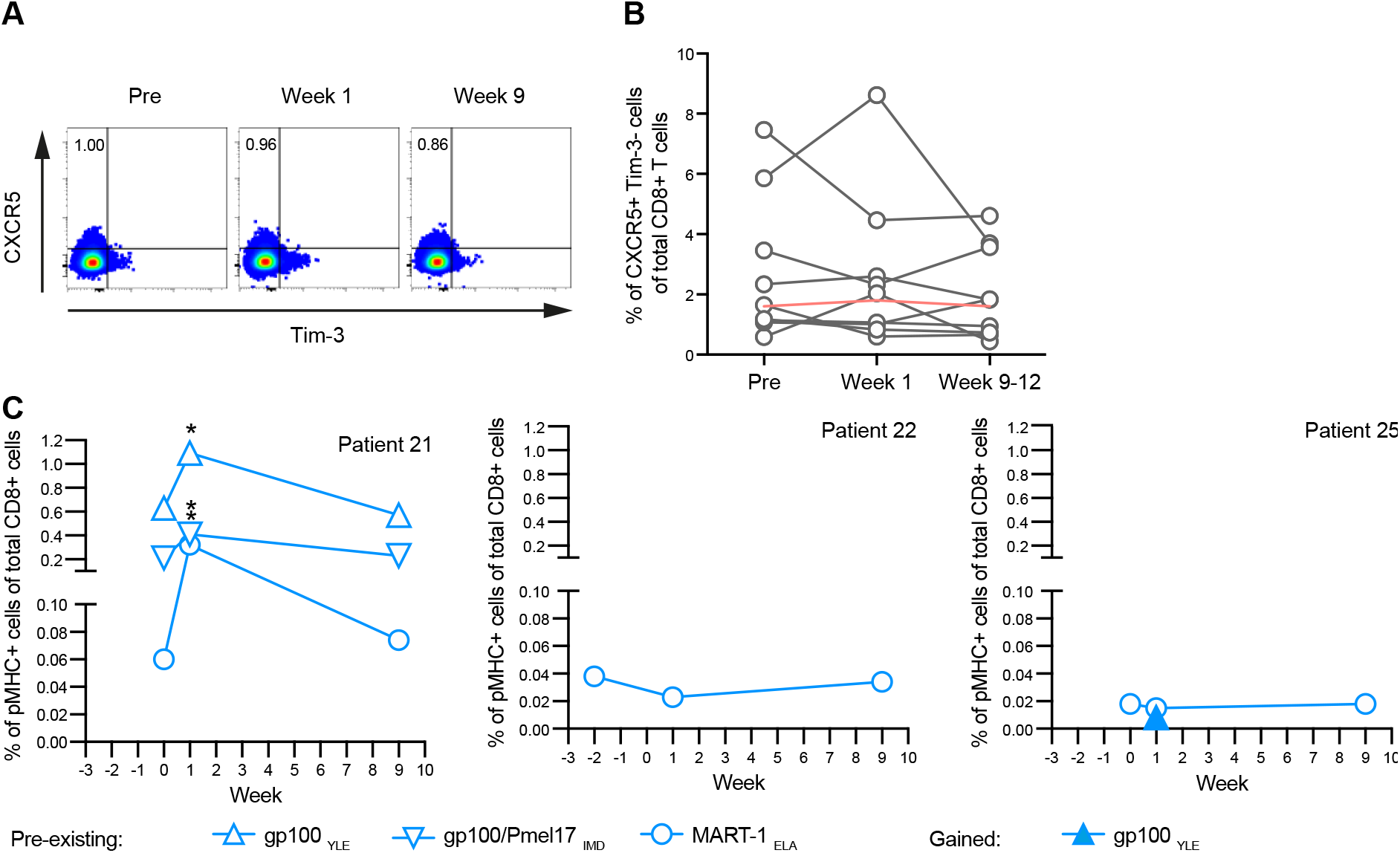
Assessment of (melanoma-reactive) CD8 T cells at week 1 after PD-1 blockade. **(A)** Representative flow cytometry plots showing the kinetics of bulk CXCR5^+^ Tim-3^-^ CD8 T cells in patient 21. The frequency is shown on the top left for each time point. **(B)** Kinetics of CXCR5^+^ Tim^-^ bulk CD8 T cells for individual patients (n=8, grey) following anti-PD-1 therapy. The median of all patients (red) was calculated for each individual time point. Statistical significance was tested with two-tailed Wilcoxon matched-pairs test, p=0.01. **(C)** Kinetics of melanoma-reactive CD8 T cell responses detected in four melanoma patients following anti-PD-1 therapy. A ≥ 2-fold increase (on vs. pre) in magnitude is indicated (*). A representative example of the full gating strategy is shown in Figure S3. pMHC: peptide-major histocompatibility complex.

### Clinical outcome of PD-1 blockade does not correlate with alterations of the melanoma-reactive CD8 T cell response

The response rate to anti-PD-1 therapy in stage IV melanoma patients is approximately 35 to 40% across multiple studies *(27)*. To understand whether our cohort was representative, we examined the clinical outcome of the patients included in this study. The objective response rate in the study cohort was 50% (Table 1). In the vast majority (10/12) of patients with clinical benefit no changes in magnitude or breadth of the circulating melanoma-reactive CD8 T cell response were observed (Table S2). Alterations of the melanoma-reactive CD8 T cell response were observed in a minority of patients, including two responders and two non-responders. These results demonstrate that the response rate to anti-PD-1 therapy in the analyzed patient cohort was comparable to previous studies *(27)* and that the effect of PD-1 blockade on the melanoma-reactive CD8 T cell response in peripheral blood does not correlate with the clinical activity of treatment.

**Table 1:** **Demographic and clinical characteristics of the study cohort treated with anti-PD-1 therapy** (*time in months between last dose of anti-CTLA-4 and first dose of anti-PD-1 therapy: median = 6, range 1-24, CR: complete response, PR: partial response, SD: stable disease, PD: progressive disease, NA: not available).

|  | Total number of patients (n = 24) |
| --- | --- |
| Median age, years (range) | 53 (31-87) |
| Gender, n (%) |  |
| Female | 10 (41%) |
| Male | 14 (59%) |
| Anti-PD-1 therapy, n (%) |  |
| Pembrolizumab | 22 (92%) |
| Nivolumab | 2 (8%) |
| CNS metastasis, n (%) |  |
| yes | 5 (21%) |
| no | 19 (79%) |
| **WHO performance status, n (%)** |  |
| NA | 6 (25%) |
| 0 | 12 (50%) |
| 1 | 6 (25%) |
| **LDH level before first dose of anti-PD-1, n (%)** |  |
| < ULN | 18 (75%) |
| 1-2 ULN | 3 (12.5%) |
| >2 ULN | 3 (12.5%) |
| **BRAF mutation, n (%)** |  |
| Yes | 8 (33%) |
| No | 16 (67%) |
| **Previous therapies, n (%)** |  |
| BRAF or BRAF/MEK inhibitors | 5 (21%) |
| Anti-CTLA-4 therapy* | 14 (58%) |
| **Best overall response (BOR) to anti-PD-1, n (%)** |  |
| NA | 1 (4%) |
| CR | 4 (17%) |
| PR | 8 (33%) |
| SD | 5 (21%) |
| PD | 6 (25%) |

### Differential effects of CTLA-4 and PD-1 blockade on the melanoma-reactive CD8 T cell response

Our current results demonstrate that PD-1 blockade does not measurably influence the melanoma-reactive CD8 T cell response in peripheral blood. In contrast, in prior work we demonstrated that anti-CTLA-4 therapy induces broadening of the melanoma-reactive CD8 T cell repertoire in peripheral blood *(21)*. To understand whether this difference in the effect of therapy on circulating CD8 T cells holds true in patient cohorts that were treated with either PD-1 or CTLA-4 blockade and analyzed concurrently using the same technology and reagents, we analyzed samples from nine melanoma patients treated with CTLA-4 blockade (clinical characteristics shown in Table S3). Peripheral blood samples were collected pre-therapy and post-therapy (approximately 12 weeks after start of treatment). In total, 13 melanoma-reactive CD8 T cell responses were identified in eight out of nine patients with a median magnitude of 0.028%, ranging from 0.006% to 0.363% (Fig. 4A and B). In concordance with our prior analyses *(21)*, the magnitude of pre-existing melanoma-reactive CD8 T cell responses was unaltered after anti-CTLA-4 therapy, with a median fold change of 0.9 (Fig. 4C). Importantly, however, and in line with prior analyses *(21)*, a broadening of the melanoma reactive CD8 T cell repertoire was observed in five out of nine melanoma patients (56%). In summary, the parallel analysis using the same technology and reagents showed a broadening of the melanoma-reactive CD8 T cell repertoire upon CTLA-4 blockade in peripheral blood, and such a broadening was observed significantly more frequent in patients treated with anti-CTLA-4 than anti-PD-1 therapy (Fig. 4D, p=0.02).

**Figure 4:**
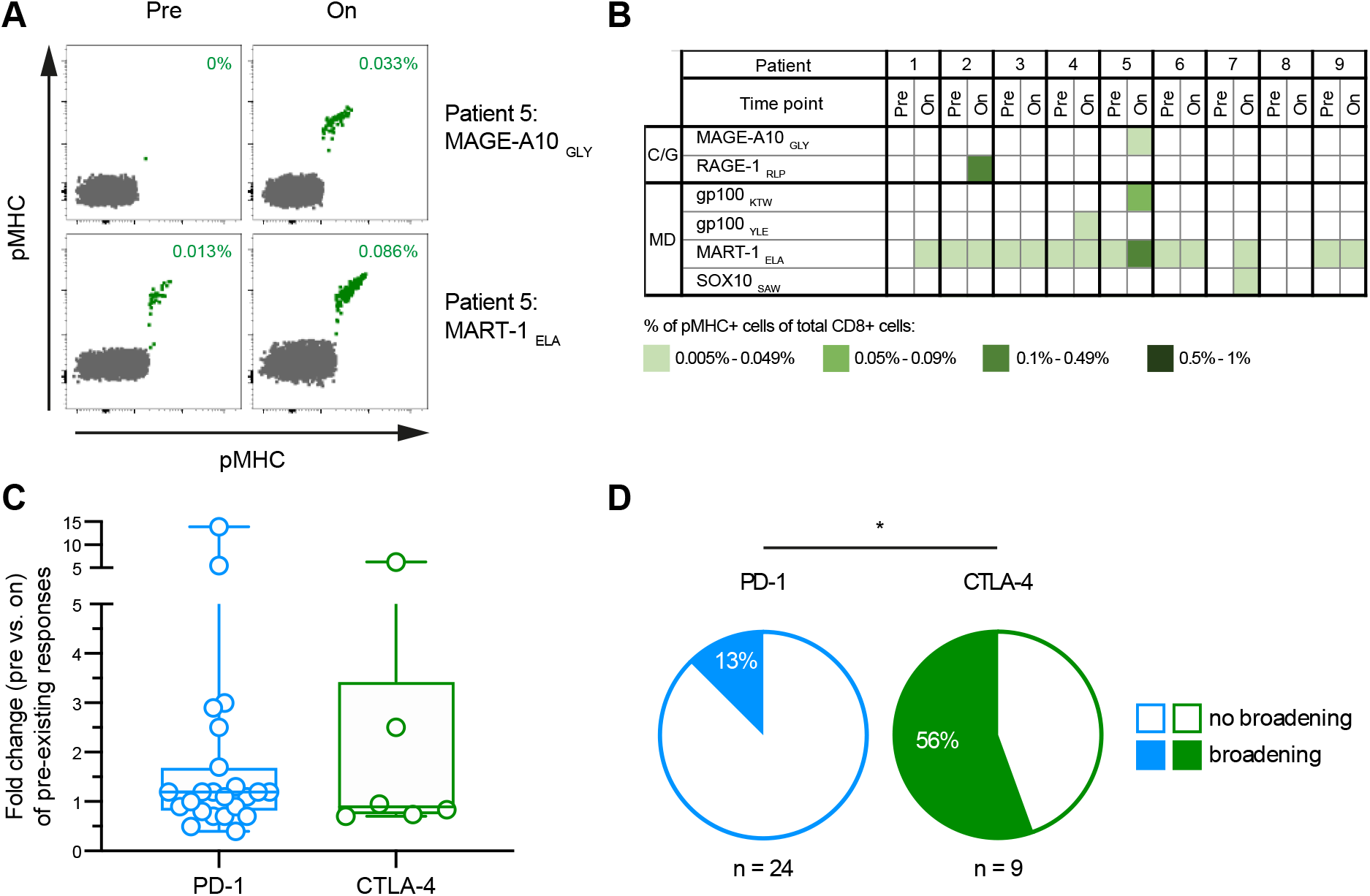
Differential effects between anti-CTLA-4 therapy and anti-PD-1 therapy on the melanoma-reactive CD8 T cell repertoire. **(A)** Representative flow cytometry plots of melanoma-reactive CD8 T cell responses before and after anti-CTLA-4 therapy from one patient. Magnitude of melanoma-reactive CD8 T cell responses represents is indicated on the top right as the percentage of total CD8 T cells for each response. Full gating strategy is provided in Figure S2. pMHC: peptide-major histocompatibility complex. **(B)** Heatmap overview of melanoma-reactive CD8 T cell responses detected in patients treated with anti-CTLA-4 therapy. C/G: cancer/germline, MD: melanocyte differentiation, pMHC: peptide-major histocompatibility complex. **(C)** Fold change in magnitude of pre-existing melanoma-reactive CD8 T cell responses detected in patients treated with anti-PD-1 or anti-CTLA-4 therapy. Boxplots represent the median and interquartile ranges, and the whiskers represent the full range. Statistical significance between the two groups was tested with two-tailed Mann-Whitney U-test, p=0.76. **(D)** Proportion of patients with 0 (no broadening) or ≥ 1 newly detectable (broadening) melanoma-reactive CD8 T cell responses after anti-PD-1 therapy (blue) or after anti-CTLA-4 therapy (green). Statistical significance between the two groups was tested with two-sided Fisher’s exact test, p=0.02.

## Discussion

The clinical success of anti-PD-1 therapy, in particular for advanced melanoma patients, has led to a major interest in understanding the mechanism of action on CD8 T cells. However, the anatomical site at which PD-1 blockade exerts its main effects remains to be fully understood with evidence for alterations of the CD8 T cell compartment at the tumor site *(5–7)* but also systemically *(14–18)*. To investigate the systemic effect of PD-1 blockade on tumor-specific CD8 T cells, we measured the impact of the treatment on the circulating melanoma-reactive CD8 T cell response directly *ex vivo* and compared these findings to the effects of CTLA-4 blockade.

Boosting or broadening of the melanoma-reactive CD8 T cell response upon PD-1 blockade was not observed in the vast majority of patients despite the objective clinical response rate of 50%. Based on these data, the previously reported proliferation of CD8 T cells subsets upon PD-1blockade *(14–17)*, may reflect proliferation of CD8 T cells with other antigen specificities. In line with this hypothesis, analysis of gp100-specific CD8 T cells of two melanoma patients showed expansion of tumor-reactive CD8 T cells upon PD-1 blockade in the tumor whereas an expansion of the T cell response was detected infrequently in peripheral blood *(16)*. Second, contrary to previous studies suggesting a potential role of PD-1 blockade in priming of novel CD8 T cell responses *(10–13)*, our data show that PD-1 blockade does not significantly increase the breadth of the melanoma-reactive CD8 T cell response. In line with these findings, a recent study has demonstrated in a mouse model that anti-PD-1 therapy as monotherapy is insufficient for the priming of naïve tumor-specific CD8 T cells *(28)*. Furthermore, our previous *(21)* and current analysis using the same technology and reagents showed that in contrast to anti-PD-1 therapy, CTLA-4 blockade increases the breadth of the melanoma-reactive CD8 T cell repertoire in peripheral blood.

The current study has the following limitations. While our analyses involve the measurement of 71 potential CD8 T cell responses, it is limited to shared melanoma antigens restricted to the HLA-A*02:01 allele. We focused on HLA-A*02:01 because the vast majority of previously identified melanoma-associated antigens are restricted to this allele. Despite our focus on only one of the potentially six different HLA alleles of each patient, melanoma-reactive CD8 T cell responses were identified in the vast majority (79%) of patients. In addition, the focus on shared melanoma antigens, rather than patient-specific neoantigens may lead us to underestimate the effects of immune checkpoint blockade. However, the observation that a broadening of the CD8 T cell response against shared melanoma antigens is observed in the current study and our previous work *(21)* provides some evidence against this possibility. While the current data argue against a profound effect of PD-1 blockade on the circulating tumor-specific CD8 T cell compartment, we observed three newly detectable CD8 T cell responses upon PD-1 blockade in a cohort of 24 patients. In addition, we have previously reported a newly detectable neoantigen-specific CD8 T cell responses upon anti-PD-1 therapy in one NSCLC patient *(29)*. Whether this represents a modest effect of PD-1 blockade on the breadth of the circulating tumor-specific CD8 T cell response will require analysis of a substantially larger patient cohort. Nevertheless, with the data available, it can be concluded that broadening of the circulating tumor-reactive CD8 T cell repertoire upon PD-1 blockade is rare compared to CTLA-4 blockade. Based on this observation and the fact that clinical activity of PD-1 blockade is substantially higher than that of CTLA-4 blockade, it may be postulated that a substantial part of the immunomodulatory effect of PD-1 blockade occurs at the tumor site, and this could include the presence of tertiary lymphoid structures at that site.

In summary, this study demonstrates that contrary to CTLA-4 blockade, PD-1 blockade does not lead to significant alterations of circulating CD8 T cells with a defined tumor specificity. In future work, it will be of interest to investigate the effect of anti-PD-1 therapy on defined tumor-reactive CD8 T cell populations at the tumor site. We note, however, that the high variability in the presence and magnitude of tumor-specific CD8 T cells even between tumor fragments of the same tumor piece *(24)* is likely to limit the ability to detect therapy-induced alterations in pre- and post-treatment biopsies. Conceivably, analysis of the cell cycle state of intratumoral tumor-reactive CD8 T cells identified using the pMHC multimer technology *(30)* may form a more sensitive approach.

## Materials and Methods

### Patient material

Peripheral blood mononuclear cell (PBMC) samples were obtained from stage IV melanoma patients undergoing immune checkpoint targeting therapy either at the Netherlands Cancer Institute (Amsterdam, The Netherlands), the University of California (Los Angeles, USA) or the University Hospital Essen (Essen, Germany). The patient cohort treated with anti-PD-1 therapy received either pembrolizumab 2 mg/kg or a fixed dose of 150-200 mg intravenously every 3 weeks, or nivolumab 3 mg/kg or in a fixed dose of 240 mg intravenously every 2 weeks in an expanded access program or according to the label after approval. The patient cohort treated with anti-CTLA-4 therapy received ipilimumab intravenously in a dose of 3 mg/kg every 3 weeks for a maximum of four cycles. Clinical response was evaluated according to response evaluation criteria in solid tumors (RECIST) version 1.1 *(31)*.

The study was conducted in accordance with the Declaration of Helsinki after approval by the institutional review boards of all centers. PBMC samples were collected in accordance with local guidelines and following signed informed consent. PBMCs were isolated using standard Ficoll gradient centrifugation separation according to local operating procedures. After isolation, PBMCs were cryopreserved in liquid nitrogen, in fetal calf serum (FCS) with 10% dimethyl sulfoxide (DMSO), or in FCS supplemented with RPMI and 10% DMSO. For the analysis of melanoma-reactive CD8 T cell responses, patients were selected based on four-digit genotyping for HLA-A*02:01.

### Epitopes

Melanoma-reactive CD8 T cell responses were identified using an HLA-A*02:01-restricted epitope panel including 71 epitopes (Table S1) derived from shared melanoma antigens. As positive control for the generation of pMHC multimers, two viral epitopes derived from Epstein-Barr virus and influenza A were included in the epitope panel (Figure S1). Peptides derived from shared melanoma-associated, viral antigens, as well as ultraviolet (UV) cleavable peptides were synthesized at the Division of Chemical Immunology, Leiden University Medical Center (Leiden, the Netherlands) as previously published *(32)*.

### Generation of pMHC multimers

MHC HLA-A*02:01 allele monomers were generated with an UV-cleavable epitope as previously published *(32)*. Specific pMHC complexes used for the identification of antigen-specific CD8 T cell responses were generated by UV-induced ligand exchange in a 96-well format as previously described *(30)*. In brief, MHC monomers loaded with UV-cleavable peptide (100 μg/ml) were exposed to 366-nm UV light (CAMAG) for 1 hour at 4 °C in the presence of a specific rescue peptide (200 μM). Subsequently, generated pMHC monomers were fluorescently labeled with streptavidin conjugates.

For data acquisition on the BD LSRII the following amounts of 10 different fluorescent streptavidin conjugates were added to 10 μl of pMHC monomer (100 μg/ml): 0.6 μl of SA-APC (Invitrogen, S868), 1.5 μl of SA-QD655 (Invitrogen, Q10121MP), 1 μl of SA-QD625 (Invitrogen, A10196), 0.4 μl of BV421 (BioLegend, 405225), 1 μl of SA-QD800 (Invitrogen, Q10171MP), 1.5 μl of SA-QD705 (Invitrogen, Q10161MP), 1.5 μl of SA-QD605 (Invitrogen, Q10101MP), 1.25 μl of SA-QD585 (Invitrogen, Q10111 MP), 1.1 μl of SA-PE-Cy7 (Invitrogen, SA1012) and 0.9 μl of SA-PE (Invitrogen, S866). For each pMHC monomer, conjugation was performed with two of these fluorochromes resulting in up to 37 dual fluorescent codes.

For data acquisition on the BD FACSymphony the following amounts of 14 different fluorescent streptavidin conjugates were added to 10 μl of pMHC monomer (100 μg/ml): 1 μl of SA-BB790 (BD, custom), 1 μl of SA-BB630 (BD, custom), 1 μl of SA-APC-R700 (BD, 565144), 0.6 μl of SA-APC (Invitrogen, S868), 1 μl of SA-BV750 (BD, custom), 2 μl of SA-BV650 (BD, 563855), 2 μl of SA-BV605 (BD, 563260), 2 μl of SA-BV480 (BD, 564876), 2 μl of SA-BV421 (BD, 563259), 1 μl of SA-BUV615 (BD, custom), 1.5 μl of SA-BUV563 (BD, 565765), 2 μl of SA-BUV395 (BD, 564176), 0.4 μl of SA-BYG670 (BD, custom) and 0.9 μl of SA-PE (Invitrogen, S866). For each pMHC monomer, conjugation was performed with two of these fluorochromes resulting in up to 74 dual fluorescent codes.

Subsequently, fluorescently labeled pMHC multimers were incubated for 30 min on ice. Finally, NaN_3_ (0.02% w/v) and an excess of D-biotin (26.4 mM, Sigma) were added to block residual binding sites. Fluorescent pMHC multimers were used for T cell staining after a minimum incubation period of 20 min at 4°C.

### Combinatorial encoding of pMHC multimers and surface marker staining

PBMCs were thawed and recovered in RPMI supplemented with human serum and desoxyribonuclease (DNAse) for one hour. Antigen-specific CD8 T cell responses were confirmed using a different dual-color code fluorochrome combination in each of the two independent experiments, the initial screen and the confirmation. All samples were analyzed on the BD LSRII. A subset of samples was analyzed on both, the BD LSRII and the BD FACSymphony, to confirm consistence between the two instruments. Analysis of Tim-3 and CXCR5 expression on CD8 T cells was only assessed in the confirmation on the BD FACSymphony due to better detection sensitivity on the BD FACSymphony as compared to BD LSRII. Before flowcytometric analysis, cells were washed twice.

For the analysis on the BD LSRII the following amount of fluorescently labeled pMHC multimers were used to stain T cells: 2 μl of SA-APC-pMHC, 0.5 μl of SA-QD655-pMHC, 1 μl of SA-QD625-pMHC, 1 μl of BV421-pMHC. 1.5 μl of SA-QD800-pMHC, 1.1 μl of SA-QD705-pMHC, 0.75 μl of SA-QD605-pMHC, 1 μl of SA-QD585-pMHC, 1.3 μl of SA-PE-Cy7-pMHC and 0.6 μl of SA-PE-pMHC. Final staining volume was 189 μl for the initial screen or 100 ul for the confirmation. Cells were incubated for 15 min in 37°C together with the pMHC multimers. Subsequently samples were stained with 2 μl of anti-CD8-AF700 (Invitrogen, MHCD0829), 1 μl of anti-CD4-FITC (BD, 345768), 1 μl of anti-CD14-FITC (BD, 345784) 1 μl of CD16-FITC (BD, 335035), 3 μl of anti-CD19-FITC (BD, 345776) and 0.5 μl of LIVE/DEAD Fixable IR Dead Cell Stain Kit (Invitrogen, L10119) and incubated on ice for 20 min.

For the analysis on the BD FACSymphony the following amount of fluorescently labeled pMHC multimers were used to stain T cells: 1 μl of SA-BB790-pMHC, SA-BB630-pMHC, SA-APC-R700-pMHC, SA-BV750-pMHC, SA-BV650-pMHC, SA-BV605-pMHC, SA-BV480-pMHC, SA-BV421-pMHC, SA-BUV615-pMHC, SA-BUV563-pMHC, SA-BUV395-pMHC, SA-BYG670-pMHC, SA-PE-pMHC and 2 μl of SA-APC-pMHC. The cells were stained using Brilliant Staining Buffer Plus (BD, 563794) according to manufacturer instructions. Final staining volume was 192 μl for the initial screen or 100 ul for the confirmation. Cells were incubated for 15 min at 37°C. Subsequently cells were stained with 2 μl of anti-CD8-BUV805 (BD, 564912), 1 μl of anti-CD4-APC-H7 (BD, 641398), 1 μl of anti-CD14-APC-H7 (BD, 560180) 1 μl of CD16-APC-H7 (BD, 560195), 3 μl of anti-CD19-APC-H7 (BD, 560252) and 0.5 μl of LIVE/DEAD Fixable IR Dead Cell Stain Kit (Invitrogen, L10119) and incubated on ice for 20 min. In the confirmation, cells were additionally stained with 1 μl of anti-CXCR5-BB515 (BD, 564625), 2 μl of anti-Tim-3-PE-Cy7 (eBioscience, 25-3109-41).

### Identification of antigen-specific CD8 T cell responses

Analysis of antigen-specific CD8 T cell responses was carried out without prior knowledge about clinical patient characteristics to avoid experimental bias. The following gating strategy was applied to identify CD8 T cells: (i) selection of live (IRDye dim) single-cell lymphocytes [forward scatter (FSC)-W/H low, side scatter (SSC)-W/H low, FSC/SSC-A], (ii) selection of anti-CD8 and ‘dump’ (anti-CD4, anti-CD14, anti-CD16, anti-CD19) negative cells. Antigen-specific CD8 T cell responses that were positive for two and only two pMHC multimer channels were identified using Boolean gating. The full gating strategy used on the BD LSRII and the BD FACSymphony is shown in Figure S2 and Figure S3, respectively. Cutoff values for the definition of positive responses were ≥ 0.005% of total CD8 T cells and ≥ 10 events in both experiments, the initial screen and the confirmation. A minimum of 50,000 CD8 T cells were acquired per sample. To reduce person-bias of manual gating, only positive responses that were confirmed by two independent people in both experiments were defined as real (Figure S2B and C). The average response magnitude that was determined by two independent people from the initial screen was used for statistical analysis. Data was analyzed using either the BD FACSDiva v.8.0.1 or the FlowJo 10.5.3 software. To monitor the reproducibility of the assay system, reference samples with up to 10 CD8 T cell responses present at varying frequencies were included in each analysis.

### Flow cytometer settings

On the BD LSRII the following 13-color instrument settings were used: blue laser (488 nm at 100 mW): FITC, 505LP, 525/50BP. Red laser (637 nm at 40 mW): APC, 670/14BP; AF700, 685LP, 710/50BP; IRDye, 750LP, 780/60BP. Violet laser (405nm at 100mW): BV421, 450/50BP; QD625, 610LP, 625/20BP; QD655, 635LP, 655/8BP. UV laser (355 nm at 20 mW): QD585, 570LP, 5812/15BP; QD605, 595LP, 605/12BP; QD705, 685LP, 710/50BP; QD800, 750LP, 780/60BP. Yellow-green laser (561 nm at 50 mW): PE, 585/15BP, PE-Cy7, 795LP, 780/60BP. On the BD FACSymphony the following 18-color instrument settings were used: blue laser (488 nm at 200 mW): BB515, 505LP, 530/30BP; BB630, 600LP, 610/20BP; BB790, 750LP, 780/60BP. Red laser (637 nm at 140 mW): APC, 670/30BP, APC-R700, 690LP, 630/45BP, IRDye and APC-H7, 750LP, 780/60BP. Violet laser (405 nm at 100 mW): BV421, 420LP, 431/28BP; BV480, 455LP, 470/20BP; BV605, 565LP, 605/40BP; BV650, 635LP, 661/11BP; BV750, 735LP, 750/30BP. UV laser (355 nm at 75 mW): BUV395, 379/28BP, BUV563, 550LP, 580/20BP; BUV615, 600LP, 615/20BP; BUV805, 770LP, 819/44BP. Yellow-green laser (561 nm at 150 mW): PE, 586/15BP; BYG670, 635LP, 670/30BP; PE-Cy7, 750LP, 780/60BP. Appropriate compensation controls were included in each analysis.

### Statistical analysis

Wilcoxon matched-pairs signed rank test was used to assess changes in the number and magnitude of antigen-specific CD8 T cell responses detected pre and on/after therapy. Differences between patient groups were assessed using the non-parametric Mann-Whitney *U*-test. Statistical significance for associations between categorical variables was determined by Fisher’s exact test. Statistical analysis was performed using PRISM 8 (Version 8.1.2).

## Acknowledgments

We thank the patients and their families for their participation in the study. We thank M. van Baalen, A. Pfauth, and F. van Diepen for their flow cytometric support. We thank R. Balderas from BD Biosciences, C. Talavera from the University of Leiden, M. Toebes, M. van den Braber from the Netherlands Cancer Institute for providing us with essential reagents for our experiments. We thank the NKI-AVL Core Facility Molecular Pathology & Biobanking for their support and for supplying us with NKI-AVL Biobank material.

## Author contributions

Conceptualization: A.G., T.N.S., J.B.A.G.H. and P.K. Data curation: A.G. Investigation: flow cytometry data was acquired by A.G., D.P., S.P. and P.K., and immunofluorescence staining data was collected by M.C.V. Formal analysis: flow cytometry data was analyzed by A.G., D.P. and P.K., and immunofluorescence staining data was analyzed by H.D. and M.C.V. Validation: A.G. and P.K. Methodology: A.G. and P.K. Resources: E.A.R., M.W.R, J.H.v.d.B, A.B., D.S., B.S., J.B.A.G.H. and C.B. provided patient material and clinical data. Project administration: A.G., D.P. and P.K. Funding acquisition: P.K. Supervision: P.K. Visualization: A.G. Writing original draft: A.G. and P.K. Reviewing and editing manuscript: all authors.

## Data and materials availability

Flow cytometry files can be found at repository (placeholder). Patient material from the University of California (Los Angeles, USA) and the University Hospital Essen (Essen, Germany) were obtained under a material transfer agreement.

