## Supplementary Material for "Differential effects of PD-1 and CTLA-4 blockade on the melanoma-reactive CD8 T cell response"

### Supplementary Materials

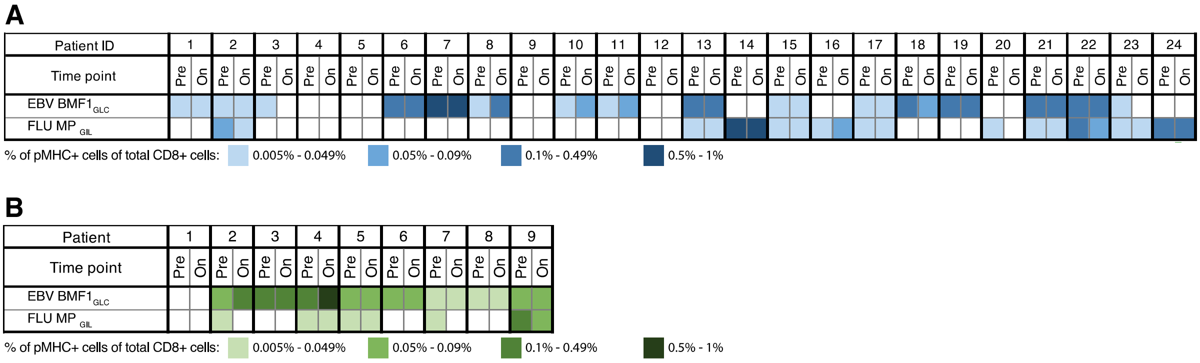


**Figure S1: Overview of detected virus-specific CD8 T cell responses (positive control for the generation of pMHC multimers).**

**(A)** Heatmap overview of virus-specific CD8 T cell responses detected in patients treated with anti-PD-1 therapy.

**(B)** Heatmap overview of virus-specific CD8 T cell responses detected in patients treated with anti-CTLA-4 therapy.

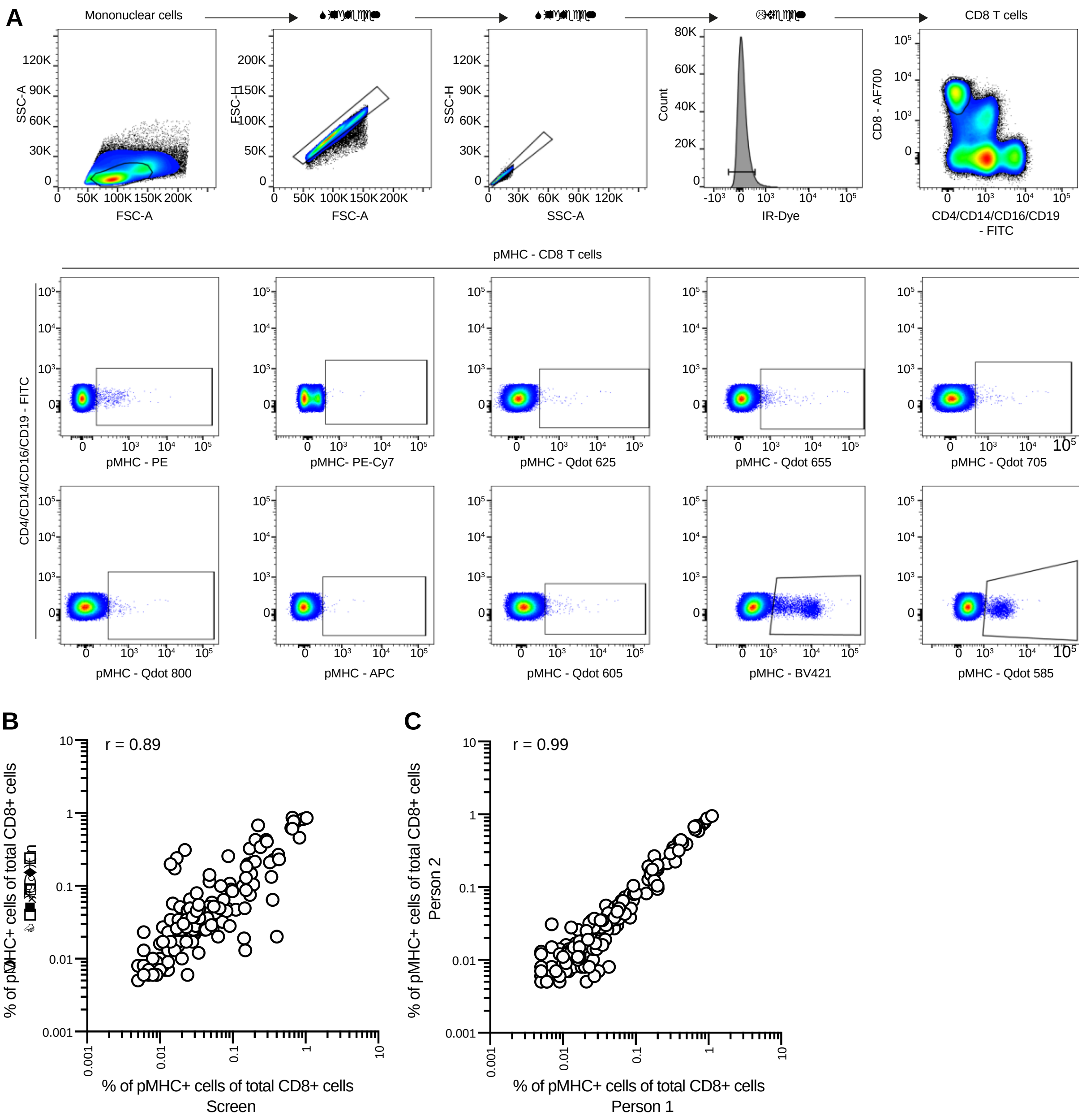


**Figure S2: Detection of antigen-specific CD8 T cell responses on the BD LSRII.**

**(A)** Representative gating strategy used for the identification of melanoma-reactive CD8 T cell responses used on the BD LSRII. Single mononuclear cells were selected by FSC and SSC. Within the single cells, live cells were gated by a live/dead stain. CD8 T cells were selected based on gating on single live mononuclear cells positive for anti-CD8-AF700 and negative for anti-CD4/-CD14/-CD16/-CD19-FITC. Melanoma-reactive CD8 T cells were identified based on selection of CD8 T cells positive for only two of the 10 fluorescent pMHC multimers using Boolean gating. pMHC: peptide-major histocompatibility complex.

**(B)** Correlation of the magnitude of antigen-specific CD8 T cell responses (% of total CD8 cells, n = 153) determined by person 1 (analysis in FlowJo 10.5.3) versus person 2 (analysis in DIVA). Data from the initial screen is shown. R value was calculated using Pearson's correlation.

**(C)** Correlation of the magnitude of antigen-specific CD8 T cell responses (% of total CD8 cells, n = 157) determined in two independent experiments (the initial screen and the confirmation). R value was calculated using Pearson's correlation.

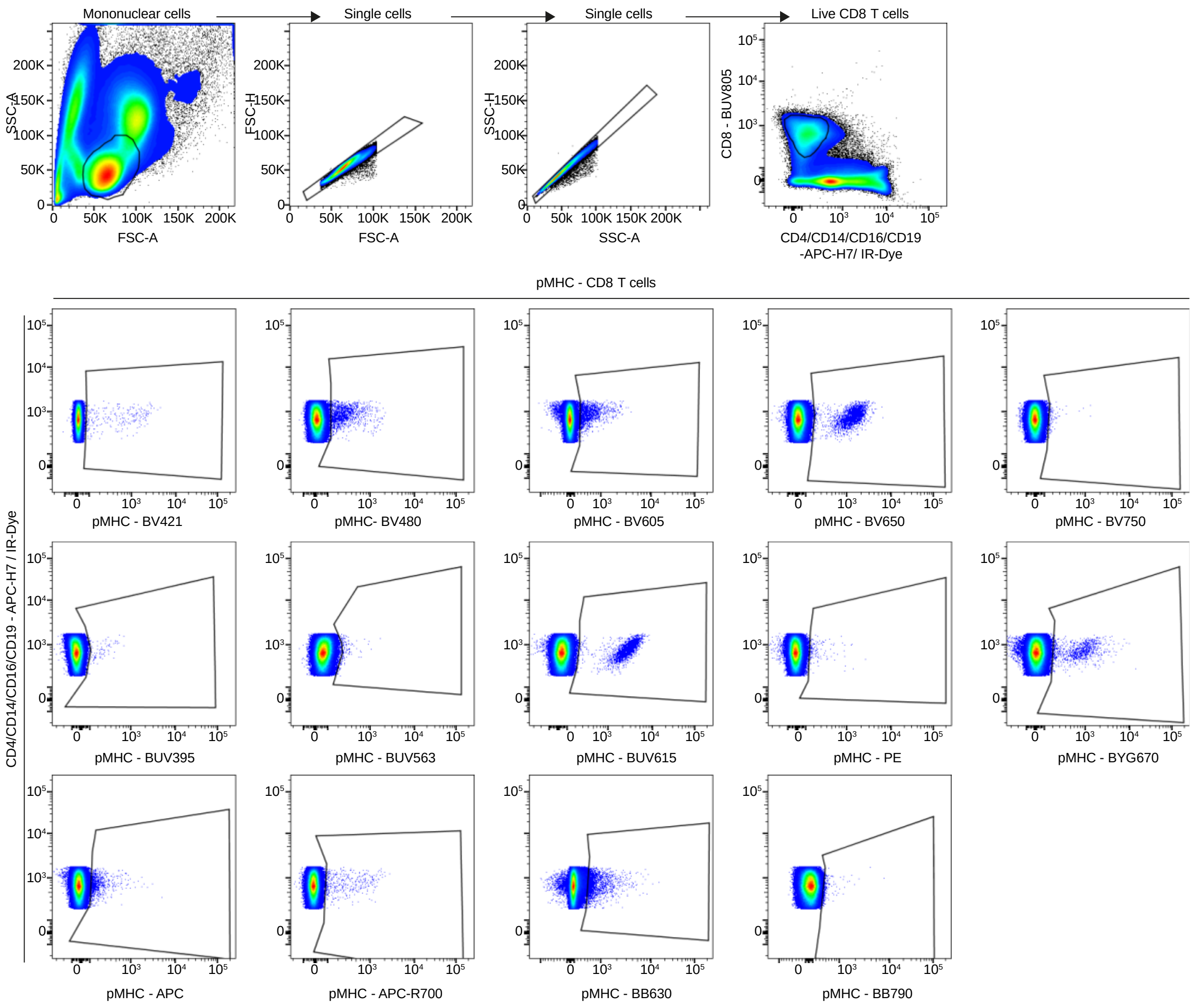


**Figure S3: Detection of antigen-specific CD8 T cell responses on the BD FACSymphony.**

Representative gating strategy used for the identification of melanoma-reactive CD8 T cell responses used on the BDFACSymphony. Single mononuclear cells were selected by FSC and SSC. Within the single cells, live CD8 T cells were selected based on gating on single live mononuclear cells positive for anti-CD8-BUV805, dim of live/dead and negative for anti-CD4/-CD14/-CD16/-CD19-APC-H7. Melanoma-reactive CD8 T cells were identified based on selection of CD8 T cells positive for only two of the 14 fluorescent pMHC multimers using Boolean gating. pMHC, peptide-major histocompatibility complex.

**Table S1: List of 71 melanoma-associated epitopes derived from different antigen sources and 2 viral epitopes including the amino acid sequences that were used for the analysis in this study.**

| Antigen | Epitope | Sequence |
|---|---|---|
| Cancer Germline (C/G) | HERV-K-MEL $_{MLA}$ | MLAVISCAV |
| Cancer Germline (C/G) | LAGE-1 $_{MLM}$ | MLMAQEALAFL |
| Cancer Germline (C/G) | LAGE-1 $_{SLL}$ | SLLMWITQC |
| Cancer Germline (C/G) | MAGE-A1 $_{KVL}$ | KVLEYVIKV |
| Cancer Germline (C/G) | MAGE-A10 $_{GLY}$ | GLYDGMEHL |
| Cancer Germline (C/G) | MAGE-A2 $_{KMV}$ | KMVELVHFL |
| Cancer Germline (C/G) | MAGE-A2 $_{LVH}$ | LVHFLLLKY |
| Cancer Germline (C/G) | MAGE-A2 $_{LVQ}$ | LVQENYLEY |
| Cancer Germline (C/G) | MAGE-A2 $_{YLQ}$ | YLQLVFGIEV |
| Cancer Germline (C/G) | MAGE-A3 $_{FLW}$ | FLWGPRALV |
| Cancer Germline (C/G) | MAGE-A3 $_{KVA}$ | KVAELVHFL |
| Cancer Germline (C/G) | MAGE-A3 $_{LVF}$ | LVFGIELMEV |
| Cancer Germline (C/G) | MAGE-A4 $_{GVY}$ | GVYDGREHTV |
| Cancer Germline (C/G) | MAGE-A6 $_{YLE}$ | YLEYRQVPV |
| Cancer Germline (C/G) | MAGE-A8 $_{GLM}$ | GLMDVQIPT |
| Cancer Germline (C/G) | MAGE-A8 $_{KVA}$ | KVAELVRFL |
| Cancer Germline (C/G) | MAGE-A9 $_{ALS}$ | ALSVMGVYV |
| Cancer Germline (C/G) | MAGE-C2 $_{ALK}$ | ALKDVEERV |
| Cancer Germline (C/G) | MAGE-C2 $_{KVL}$ | KVLEFLAKL |
| Cancer Germline (C/G) | MAGE-C2 $_{LLF}$ | LLFGLALIEV |
| Cancer Germline (C/G) | MAGE-C2 $_{TLD}$ | TLDEKVAELV |
| Cancer Germline (C/G) | MAGE-C2 $_{VIW}$ | VIWEVLNAV |
| Cancer Germline (C/G) | NY-ESO1 $_{QLS}$ | QLSLLMWIT |
| Cancer Germline (C/G) | NY-ESO1 $_{SLL}$ | SLLMWITQCFL |
| Cancer Germline (C/G) | NY-ESO1 $_{SLL}$ | SLLMWITQA |
| Cancer Germline (C/G) | PRAME $_{ALY}$ | ALYVDSLFFL |

| | | |
|---|---|---|
| Cancer Germline (C/G) | PRAME $_{SLL}$ | SLLQHLIGL |
| Cancer Germline (C/G) | PRAME $_{SLY}$ | SLYSFPEPEA |
| Cancer Germline (C/G) | PRAME $_{VLD}$ | VLDGLDVLL |
| Cancer Germline (C/G) | RAGE-1 $_{LKL}$ | LKLSGVVRL |
| Cancer Germline (C/G) | RAGE-1 $_{PLP}$ | PLPPARNGGL |
| Cancer Germline (C/G) | SSX-2 $_{KAS}$ | KASEKIFYV |
| Cancer Germline (C/G) | SSX-2 $_{RLQ}$ | RLQGISPKI |
| Cancer Germline (C/G) | TAG-1 $_{SLG}$ | SLGWLFLLL |
| Cancer Germline (C/G) | TRAG-3 $_{GLI}$ | GLIQLVEGV |
| Cancer Germline (C/G) | TRAG-3 $_{ILL}$ | ILLRDAGLV |
| Cancer Germline (C/G) | TRAG-3 $_{VLG}$ | VLGEAWRDQV |
| Cancer Germline (CG) | CDCA1/NUF2 $_{YMM}$ | YMMPVNSEV |
| Cancer Germline (CG) | CDCA1/NUF3 $_{KLA}$ | KLATAQFKI |
| Cancer Germline (CG) | CML28 (EXOSC5) $_{ALV}$ | ALVDAGVPM |
| Cancer Germline (CG) | DAM-6, -10 (MAGE-B1, -B2) $_{FLW}$ | FLWGPRAYA |
| Melanocyte differentiation (MD) | gp100 / Pmel17 $_{AML}$ | AMLGTHTMEV |
| Melanocyte differentiation (MD) | gp100 $_{KTW}$ | KTWGQYWQV |
| Melanocyte differentiation (MD) | gp100 $_{LLD}$ | LLDGTATLRL |
| Melanocyte differentiation (MD) | gp100 $_{MLG}$ | MLGTHTMEV |
| Melanocyte differentiation (MD) | gp100 $_{RLM}$ | RLMKQDFSV |
| Melanocyte differentiation (MD) | gp100 $_{RLP}$ | RLPRIFCSC |
| Melanocyte differentiation (MD) | gp100 $_{SLA}$ | SLADTNSLAV |
| Melanocyte differentiation (MD) | gp100 $_{VLY}$ | VLYRYGSFSV |
| Melanocyte differentiation (MD) | gp100 $_{YLE}$ | YLEPGPVTA |
| Melanocyte differentiation (MD) | gp100/Pmel17 $_{IMD}$ | IMDQVPFSV |
| Melanocyte differentiation (MD) | Melan-A / MART-1 $_{ELA}$ | ELAGIGILTV |
| Melanocyte differentiation (MD) | Melan-A / MART-1 $_{ILT}$ | ILTVILGVL |
| Melanocyte differentiation (MD) | NY-MEL-1 $_{VLH}$ | VLHWDPETV |
| Melanocyte differentiation (MD) | SOX10 $_{AWI}$ | AWISKPPGV |
| Melanocyte differentiation (MD) | SOX10 $_{SAW}$ | SAWISKPPGV |
| Melanocyte differentiation (MD) | TRP-2 $_{FVW}$ | FVWLHYYSV |
| Melanocyte differentiation (MD) | TRP-2 $_{SLD}$ | SLDDYNHLV |
| Melanocyte differentiation (MD) | TRP-2 $_{SVY}$ | SVYDFFVWL |
| Melanocyte differentiation (MD) | TRP-2 $_{TLD}$ | TLDSQVMSL |
| Melanocyte differentiation (MD) | TRP-2 $_{VYD}$ | VYDFFVWLHY |
| Melanocyte differentiation (MD) | Tyrosinase $_{CLL}$ | CLLWSFQTSA |
| Melanocyte differentiation (MD) | Tyrosinase $_{MLL}$ | MLLAVLYCL |
| Melanocyte differentiation (MD) | Tyrosinase $_{YMD}$ | YMDGTMSQV |
| Overexpressed (OE) | B-RAF $_{LAT}$ | LATEKSRWS |
| Overexpressed (OE) | BING-4 (WDR46) $_{CQW}$ | CQWGRLWQL |
| Overexpressed (OE) | CDK4 $_{ACD}$ | ACDPHSGHFV |
| Overexpressed (OE) | GnTV $_{VLP}$ | VLPDVFIRCV |
| Overexpressed (OE) | Meloe-1 $_{TLN}$ | TLNDECWPA |
| Overexpressed (OE) | Meloe-2 $_{RLP}$ | RLPPKPPLA |
| Overexpressed (OE) | PRDX5 (OMT3-12) $_{AMA}$ | AMAPIKVRL |
| Viral (V) | EBV BMF1 $_{GLC}$ | GLCTLVAML |
| Viral (V) | FLU MP $_{GIL}$ | GILGFVFTL |

**Table S2: Overview of the clinical outcome and the systemic effects detected approx. 12 weeks post PD-1 blockade for each patient.** (*Responses: melanoma-reactive CD8 T cell response, CR: complete response, PR: partial response, SD: stable disease, PD: progressive disease, NA: not available).

| Patient | Best overall response | *Responses detected pre-therapy, n | *Responses detected post-therapy, n | Broadening: novel *responses post-therapy, no / yes (n) | Boosting: fold change of pre-existing *responses post-therapy, no (n) / yes (n) |
|---|---|---|---|---|---|
| 1 | SD | 2 | 2 | no | yes (5.4) |
| 2 | PD | 1 | 1 | no | no (0.9) |
| 3 | SD | 2 | 2 | no | no (0.7) |
| 4 | NA | 1 | 3 | yes (2) | yes (13.9) |
| 5 | PD | 1 | 1 | no | no (1.3) |
| 6 | CR | 0 | 0 | no | - |
| 7 | SD | 1 | 1 | no | no (1.2) |
| 8 | PD | 1 | 1 | no | no (0.5) |
| 9 | SD | 1 | 1 | no | no (1.6) |
| 10 | PD | 1 | 2 | yes (1) | yes (2.9) |
| 11 | CR | 1 | 1 | no | no (1.2) |
| 12 | PR | 1 | 1 | no | yes (2.4) |
| 13 | PD | 1 | 1 | no | 1.2 |
| 14 | PR | 1 | 1 | no | 1.2 |
| 15 | CR | 0 | 0 | no | - |
| 16 | PR | 0 | 0 | no | - |
| 17 | CR | 1 | 1 | no | 0.8 |
| 18 | PD | 1 | 1 | no | 0.6 |
| 19 | PD | 1 | 1 | no | 0.4 |
| 20 | PR | 0 | 0 | no | - |
| 21 | PR | 3 | 3 | no | 1.1 |
| 22 | PR | 1 | 1 | no | 0.8 |
| 23 | PR | 0 | 0 | no | - |
| 24 | PR | 1 | 2 | yes (1) | 1.2 |

**Table S3: Demographic and clinical charateristics of the study cohort treated with anti-CTLA-4 therapy** (*time in months between last dose of Nivolumab and first dose of CTLA-4 blockade: median = 2, range 1-3, CR: complete response, PR: partial response, SD: stable disease, PD: progressive disease).

| | **Total number of patients (n = 9)** |
|---|---|
| **Median age, years (range)** | 64 (49-67) |
| **Gender, n (%)** | |
| Female | 4 (44%) |
| Male | 5 (56%) |
| **CNS metastasis, n (%)** | |
| yes | 1 (11%) |
| no | 8 (89%) |
| **WHO performance status, n (%)** | |
| 0 | 6 (67%) |
| 1 | 3 (33%) |
| **LDH level before first dose of anti-PD-1, n (%)** | |
| < ULN | 7 (78%) |
| 1-2 ULN | 2 (22%) |
| **BRAF mutation, n (%)** | |
| Yes | 3 (33%) |
| No | 6 (67%) |
| **Previous therapies, n (%)** | |
| BRAF or BRAF/MEK | 2 (22%) |
| Nivolumab* | 3 (33%) |
| **Best overall response (BOR) to anti-CTLA-4 therapy** | |
| CR | 0 (0%) |
| PR | 1 (11%) |
| SD | 0 (0%) |
| PD | 8 (89%) |
